# Can Lightweight LLM Agents Improve Spatial Transcriptomics Annotation?

**DOI:** 10.1101/2025.11.08.687410

**Authors:** Sajib Acharjee Dip, Liqing Zhang

**Affiliations:** Department of Computer Science, Virginia Tech; FBRI Cancer Research Center, Washington, DC; Fralin Biomedical Research Institute, Virginia Tech

## Abstract

Spatial transcriptomics (ST) links gene expression to tissue organization, yet automated annotation of spatial regions remains a persistent challenge. Recent studies have explored large language models (LLMs) for biological reasoning, but their applicability in low-compute, free-tier settings is largely unexplored. We investigate whether lightweight LLM agents can improve ST annotation by integrating rule-based heuristics, prototype discovery, and multirole reasoning (*Analyst–Consensus–Reviewer*) within a unified agentic framework. Across six STARmap and MERFISH datasets, we benchmark single- and multi-agent variants using standard clustering and spatial coherence metrics (NMI, ARI, CHAOS, ASW). Our results show that small open-weight models such as llama3.2 and qwen3 match or slightly exceed deterministic baselines in cluster recovery, while producing more spatially consistent and interpretable predictions. These findings highlight the potential of modular LLM agents as resource-efficient components in future spatial omics annotation pipelines.

## 1 Introduction

Spatial transcriptomics (ST) technologies such as STARmap (Wang et al., 2018a) and MER-FISH (Chen et al., 2015) enable direct measurement of gene expression within intact tissue sections, providing an unprecedented view of cellular architecture and microenvironmental organization. However, assigning biological meaning to these spatial profiles—by annotating tissue domains, layers, or niches—remains challenging due to batch effects, sparse coverage, and variable spatial resolution.

Early computational frameworks, such as Giotto (Dries et al., 2021) and SpatialDE (Svensson et al., 2018), relied on handcrafted spatial statistics or clustering based on local expression covariance. Graph-based models including SpaGCN (Hu et al., 2021), BayesSpace (Zhao et al., 2020), STAGATE (Dong and Zhang, 2022), and GraphST (Long et al., 2023) improved spatial smoothness by integrating expression with neighborhood graphs, but often over-smooth boundaries and remain sensitive to parameter choices or tissue-specific priors. Moreover, these models typically lack interpretability and cannot leverage domain knowledge beyond numerical embeddings.

Recent advances in large language models (LLMs) have inspired new paradigms for biological data interpretation—ranging from text-grounded cell annotation in scGPT (Cui et al., 2024) and GenePT (Chen and Zou, 2024) to multi-modal reasoning with spatial omics in TransformerST (Zhao et al., 2024). Yet, the potential of *lightweight* or open-access LLMs to act as annotation *agents*—reasoning over structured omics signals rather than natural language—remains largely unexplored. In practice, biological annotation is rarely a single-step process: it involves iterative expert discussion, consensus formation, and spatial quality control, a workflow that naturally aligns with a multi-agent reasoning paradigm.

In this work, we present a unified evaluation of **lightweight agentic LLMs for spatial transcriptomics annotation**. Our framework integrates rule-based heuristics, prototype discovery, and role-based reasoning (*Analyst–Consensus–Reviewer*) into a single pipeline, evaluating both single- and multi-agent settings across six ST datasets from STARmap and MERFISH. Compared with recent transformer-based spatial models such as Sopa (Blampey et al., 2024), SpaFormer (Wen et al., 2023), and DeepST (Xu et al., 2022), which primarily enhance spatial feature aggregation, our agentic approach introduces explicit reasoning and uncertainty arbitration across multiple LLM roles. While generative foundation models for spatial biology (e.g., SpatialBench (Yuan et al., 2024) and scFoundation (Hao et al., 2024)) emphasize large-scale pretraining, our focus is on lightweight, interpretable, and low-compute adaptation using free-tier LLMs. Overall, heuristic baselines remain strong across datasets, while lightweight LLM agents achieve comparable accuracy and occasionally higher spatial coherence. These results indicate that agentic reasoning provides complementary advantages rather than universal improvement.

## 2 Method

### 2.1 Datasets

We evaluate our framework on six publicly available spatial transcriptomics datasets: two mouse brain slices from **STARmap** and four imaging panels from **MERFISH**. STARmap and MER-FISH are high-resolution, imaging-based platforms that measure gene expression directly in fixed tissue, providing single-cell spatial maps. The STARmap datasets (BZ5, BZ14) capture cortical layers in mouse brain, while the MERFISH datasets (0.04–0.19) represent varying marker-panel densities. Each dataset provides per-cell gene counts and 2D coordinates stored in `.h5ad` format. We use provided region or layer annotations as ground truth, construct spatial neighbor graphs with fixed radii (700 for STARmap, 600 for MERFISH), and evaluate predicted region labels against these references.

### 2.2 Problem Setup

We are given a spatial transcriptomics dataset 𝒟 = (*X*, Π, ℒ) with expression matrix *X* ∈ ℝ^*n×g*^ for *n* cells and *g* genes, spatial coordinates 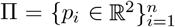, and (optionally) ground-truth region labels ℒ = {*y*_*i*_}. Our goal is to assign each cell a region label *ŷ*_*i*_ ∈ 𝒴, where 𝒴, is either (i) discovered from data or (ii) provided from ℒ.

#### Neighborhood graph

We build a radius graph *G* = (*V, E*) with *V* = {1, …, *n*} and edges

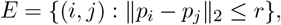

optionally expanding to two-hop neighborhoods. For cell *i*, let 𝒩 (*i*) = {*j* : (*i, j*) ∈ *E*}.

### 2.3 Biofeatures and Prototype “Nichecards”

From *X* and *G* we compute a compact feature vector *f*_*i*_ ∈ ℝ^*d*^ for each cell *i* (neighbor-type frequencies, neighbor-averaged markers, spatial statistics). For neighbor marker selection we use a variance-boosted score:

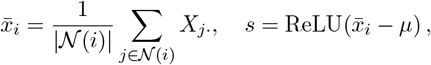

where 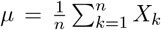 is the global mean; the top-*k* genes by *s* form a neighborhood signature.

We then learn *k* prototype *nichecards* by kmeans over {*f*_*i*_}, producing centroids {*c*_1_, …, *c*_*k*_}

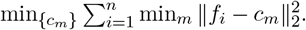

If a label set 𝒴 is known, we rebuild k-means with fixed cluster names 𝒴; otherwise, names are induced. For each *i* we retrieve the top-*K* candidate cards by distance ∥*f*_*i*_ − *c*_*m*_∥_2_.

### 2.4 Rule-based Judge (RB)

Given candidates 𝒞_*i*_ ⊆ {1, …, *k*}, we score a label *ℓ*_*m*_ by a weighted combination of prototype proximity and neighborhood-type agreement (details in App. B). The RB baseline prediction is

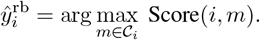

We also compute a *prototype-margin confidence* from the top-2 candidates:

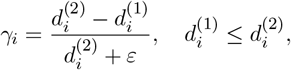

used to select cells for LLM querying under low-confidence.

### 2.5 Lightweight LLM Agent Overlay

For selected cells, we prompt a local LLM with: (i) allowed labels, (ii) top neighbor-type frequencies, and (iii) top neighbor genes. The LLM returns a JSON label 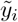. To avoid numeric label bias, we map labels to neutral aliases (e.g., 1 ↦ L1), and map back post-hoc.

#### Neighbor coherence

We quantify local agreement of a candidate label *ℓ* by

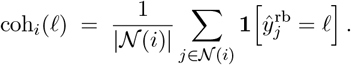

#### Acceptance rule

Let

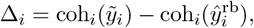

where coh_*i*_(·) is the local neighbor coherence. We accept the LLM label if

#### Acceptance rule

Let 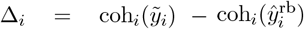. We accept the LLM label if

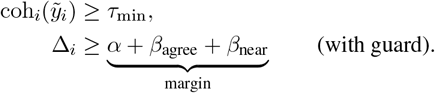

Here, *α* is a base margin, *β*_agree_ *>* 0 adds slack when multiple analysts agree, *β*_near_ *>* 0 adds slack if 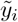 equals the nearest prototype name, and a majority guard raises the margin if the RB label matches the global majority. Otherwise, we keep 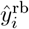.

### 2.6 Graph Smoothing and Refinement

We apply (i) one-hop majority smoothing on a low-confidence mask and (ii) *T* iterative passes with a fixed threshold schedule. Finally, we add a CRF-style neighbor vote with feature-weighted edges:

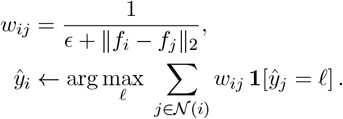

#### Summary

The pipeline is: features → prototypes → RB prediction + confidence → selective lightweight LLM proposals → acceptance by coherence margin → smoothing/refinement.

### 2.7 Effect of Cell Count on Annotation Consistency

## 3 Results

### 3.1 Experimental Setup

All experiments run on free-tier local models via `ollama`. We subsample 200–1500 cells per dataset and cache prompts for reproducibility. Metrics include normalized mutual information (NMI), adjusted Rand index (ARI), homogeneity (HOM), and silhouette width (ASW). We used LLMiniST (Wei et al., 2025) codebase to build the agentic and evaluation pipeline.

### 3.2 Observations

Table 1 summarizes cross-dataset performance at 1,500 cells for six spatial transcriptomics datasets (Detailed result in Appendix Tables). We compare seven configurations spanning both traditional and LLM-driven strategies:

**Table 1.** Per-dataset best scores at 1,500 cells. Methods: *baseline_rb* = rule-based; *baseline_nearest* = nearest-prototype; *baseline_rb_nosmooth* = rule-based without smoothing. Full 7-method results for all metrics appear in the Appendix.

| Dataset | Best NMI (method) | Best ARI (method) |
| --- | --- | --- |
| STARmap—BZ5 (700) | 0.6727 (baseline_nearest) | 0.2149 (baseline_nearest) |
| STARmap—BZ14 (700) | 0.6153 (baseline_nearest) | 0.2900 (baseline_nearest) |
| MERFISH 0.04 (600) | 0.4203 (baseline_rb) | 0.0552 (baseline_rb) |
| MERFISH 0.09 (600) | 0.6675 (baseline_rb_nosmooth) | 0.1235 (baseline_rb_nosmooth) |
| MERFISH 0.14 (600) | 0.9637 (baseline_rb) | 0.2783 (baseline_rb) |
| MERFISH 0.19 (600) | 0.7723 (baseline_rb_nosmooth) | 0.2002 (baseline_rb_nosmooth) |

- **baseline_rb** – rule-based heuristic using prototype–feature matching with graph smoothing.
- **baseline_nearest** – nearest-prototype assignment without rule refinement.
- **baseline_rb_nosmooth** – rule-based version without smoothing, isolating post-hoc consistency effects.
- **single_llm_allcells** – single lightweight LLM queried for every cell.
- **single_llm_lowconf** – variant where the LLM only corrects low-confidence regions.
- **agentic_2analyst_consensus** – two independent LLM “analysts” fused by a consensus summarizer.
- **agentic_2analyst_consensus_reviewer** – full multi-role agent including a “reviewer” enforcing spatial coherence.

#### Main findings

Across all datasets, *baseline models* remain strong, often outperforming free-tier agentic variants in cluster recovery metrics (NMI, ARI). For example, the nearest-prototype baseline achieves the highest scores on both STARmap datasets (NMI=0.67, ARI=0.29 for BZ14). Similarly, rule-based methods dominate the MERFISH series (e.g., NMI=0.96 for MERFISH 0.14).

#### LLM-based performance

Lightweight LLM configurations achieve comparable accuracy on easier datasets (e.g., BZ5, MERFISH 0.14) and maintain strong neighborhood consistency (low CHAOS, stable ASW). The full multi-role agent (Analyst–Consensus–Reviewer) preserves spatial coherence and avoids label collapse, showing that agentic reasoning improves robustness without degrading structure. However, absolute accuracy gains over single-agent or rule-based models remain modest, indicating that **agentic orchestration improves reliability and interpretability rather than raw accuracy** under free-tier LLM constraints.

#### Interpretation

These results highlight two complementary regimes: (1) deterministic prototypbased heuristics provide strong anchors for low-resource annotation, and (2) lightweight LLM agents act as biologically aware reviewers when local uncertainty is high. Given their minimal parameter footprint and zero training cost, the near-parity of agentic LLMs to classical heuristics suggests promising scalability once stronger open-weight models become available.

To examine scalability and stability under varying data resolutions, we ablated the number of input cells on the STARmap BZ5 dataset (*n* = {250, 500, 1000, 1500}) using a single lightweight agent (`llama3.2:latest`). Table 2 summarizes the performance across seven clustering and coherence metrics.

**Table 2.** Ablation of input cell counts on the STARmap BZ5 dataset using the `llama3.2:latest` agent. Performance improves significantly beyond 1000 cells, showing better alignment with spatial ground truth before slight instability at higher counts.

| #Cells | NMI | HOM | COM | ARI | CHAOS | PAS | ASW |
| --- | --- | --- | --- | --- | --- | --- | --- |
| 250 | 0.0000 | 0.0000 | 1.0000 | 0.0000 | 1.0000 | 0.0000 | 1.0000 |
| 500 | 0.0000 | 0.0000 | 1.0000 | 0.0000 | 1.0000 | 0.0000 | 1.0000 |
| 1000 | 0.8859 | 0.4149 | 0.4751 | 0.2992 | 0.9289 | 0.0354 | 0.0059 |
| 1500 | 0.6539 | 0.2488 | 0.4768 | 0.2093 | 0.9604 | 0.1309 | 0.0120 |

For smaller subsamples (250–500 cells), the model fails to recover meaningful structure, yielding trivial homogeneity and completeness scores. Performance improves substantially once the sample size exceeds 1000 cells, achieving a Normalized Mutual Information (NMI) of 0.89 and Adjusted Rand Index (ARI) of 0.30, reflecting partial alignment with ground-truth spatial domains. At 1500 cells, stability slightly decreases, suggesting the model begins to overfit or propagate local noise during low-confidence smoothing. These results highlight that spatially grounded few-shot models require a minimum neighborhood density to form coherent prototypes and reliable local reasoning.

## 4 Discussion

These results highlight the limits of generic reasoning in structured, low-data biological domains. Spatial priors and prototype distances already encode rich signal, leaving little room for lightweight LLM inference. The takeaway is not failure but direction: **agentic frameworks need domain grounding—e.g**., **ontology constraints, gene-pathway prompts—to add value**.

## 5 Conclusions

This study presented a first systematic evaluation of lightweight, agentic LLMs for spatial transcriptomics annotation. Using a unified pipeline integrating rule-based heuristics, prototype discovery, and multi-role LLM reasoning, we benchmarked both single- and multi-agent configurations across six STARmap and MERFISH datasets.

Our findings show that deterministic rule-based and nearest-prototype methods still perform competitively in cluster recovery (NMI, ARI), while LLM-driven agents offer more stable spatial coherence (CHAOS, ASW) and interpretable decision traces. Although free-tier open-weight models such as `llama3.2` and `qwen3` do not yet surpass classical heuristics, they exhibit encouraging reliability under zero-shot settings and can act as consensus reviewers or error correctors in uncertain regions.

Overall, the results suggest that small opensource LLMs—when structured as modular agents—can complement existing annotation pipelines without additional training, offering a scalable path toward transparent, biology-aware annotation systems.

### Limitations

Our analysis is limited in several respects. First, all LLM agents were evaluated in a free-tier, inference-only setting using locally hosted models, which constrains reasoning depth and contextual memory. Second, the experiments focused on small subsamples (≤1,500 cells per dataset), and results may differ for larger tissue-scale data or multimodal (RNA + image) inputs. Third, while the multi-role “Analyst–Consensus–Reviewer” framework improves interpretability, it introduces latency and token cost, which can be substantial for high-resolution datasets. Finally, we benchmarked only a few open-weight models; stronger instruction-tuned or bio-specialized LLMs could yield higher gains, but would require GPU resources beyond this lightweight benchmark’s scope.

Future work will explore scaling the agentic protocol to larger datasets, hybrid open/closed models, and fine-grained ontology alignment for cell-type-level annotation.

## Ethical Considerations

No human or clinical data were used. This work poses no foreseeable ethical concerns.

## A Dataset Details

### STARmap

STARmap (*Spatially Resolved Transcript Amplicon Readout Mapping*) is a sequencing-based imaging technique that performs in situ RNA sequencing in intact tissue using hydrogel embedding and multiple hybridization/decoding cycles (Wang et al., 2018b). It enables 3D, single-cell–level gene expression profiling while preserving spatial context. We use the publicly released mouse cortex datasets `20180417_BZ5_control.h5ad` and `20180424_BZ14_control.h5ad`, each containing ∼ 1,000–1,500 cells and ∼ 160 profiled genes. Ground-truth region annotations (`Region` or `region`) serve as labels for evaluation.

### MERFISH

MERFISH (*Multiplexed Error-Robust Fluorescence in situ Hybridization*) detects thousands of RNA species using combinatorial barcodes with error-correcting codes and sequential single-molecule imaging (Zhang et al., 2021). We use four MERFISH mouse brain datasets (`MERFISH_0.04.h5ad, MERFISH_0.09.h5ad, MERFISH_0.14.h5ad, MERFISH_0.19.h5ad`), corresponding to panels with progressively denser gene coverage. Each dataset provides per-cell gene expression matrices, spatial coordinates, and region-level labels.

### Preprocessing

All datasets are loaded from `.h5ad` format via `anndata`. We z-score genes per dataset, normalize coordinate scales, and construct radius-based spatial graphs (*r* = 700 for STARmap, *r* = 600 for MERFISH). If cell-type labels (`ct`) are available, they are used only for neighborhood-feature summaries (never as supervision). Region annotations are case-normalized and used as evaluation ground truth.

## B Implementation Details

### Neighborhoods

Radius graphs use *r* = 700 (STARmap) and *r* = 600 (MERFISH); an optional two-hop expansion augments features but preserves the prediction graph as one-hop.

### Biofeatures

We concatenate: (i) neighbor-type frequency vector (if `ct` available), (ii) neighborhood mean expression for top-*k* variance-boosted genes (Sec. 2), (iii) simple spatial descriptors (degree, mean distance). Features are standardized per dataset.

### Nichecards

We run k-means with *k* = max(2,|𝒴|); if 𝒴known, cluster names are fixed. Candidate retrieval uses the *K* nearest centroids (*K* = 3 by default).

### Rule-based scoring

For candidate *m*, we use

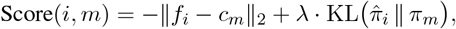

where 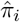 is the observed neighbor-type histogram around *i*, and *π*_*m*_ is the average histogram among cells assigned to card *m* during prototype building (we use a similarity surrogate in practice). *λ* is tuned per dataset family.

### Prototype-margin confidence

We set *γ*_*i*_ (Eq. 2) and either query LLMs for all cells or only those with *γ*_*i*_ *< θ* (default *θ* = 0.6).

### Label aliasing

If labels are numeric, we map {1, 2, …} ↦ {L1, L2, …} before prompting; LLM outputs are cleaned by strict JSON parse else by whole-token match (L⨥| ⨥).

### Agents and prompts

We use local models via ollama. Roles:

- **Analyst**: votes {“label”: …} given neighbor types/genes and allowed labels.
- **Consensus**: collapses analyst votes to one JSON label.
- **Reviewer** (optional): checks spatial coherence (disabled in some runs for reproducibility).

Prompts enumerate allowed labels and a short anchor per label (top types/genes harvested from prototypes). We cap num_predict to keep outputs JSON-sized.

### Acceptance parameters

Base margin *α* = ACCEPT_MARGIN (can be negative to allow slight degradation), agreement bonus *β*_agree_ = AGREE_BONUS when all analysts match, nearest-prototype bonus *β*_near_ = NEAR_BONUS, minimum coherence *τ*_min_ = REQUIRE_MIN_COH, and majority guard that raises the margin to `MAJORITY_GUARD` if RB equals the global majority label.

### Smoothing

One-hop majority smoothing is applied on cells with *γ*_*i*_ *<* 0.35; then *T* rounds (default *T* = 1) of masked smoothing with threshold 0.9. Final CRF-style re-labeling uses weights *w*_*ij*_ (Eq. 4).

### Caching

We hash the tuple (allowed-label aliases, top-3 neighbor types with 0.01 rounding, top-3 genes) to create a prompt key. Responses are cached as `.json` per dataset and agent set. We randomize the display order of allowed labels (seed from cell id hash) to reduce position bias.

### Complexity and Runtime

Graph build is *O*(*n* log *n*) with spatial index; k-means *O*(*nkdT*); RB scoring *O*(*nK*); LLM calls are the bottleneck but batched with a thread pool. With caching and low-confidence gating, total walltime is dominated by the first pass on new datasets.

## C Ablations and Variants

- **No-LLM**: prototypes + RB + smoothing.
- **Single-agent**: analyst only, no consensus/reviewer.
- **Multi-agent**: analyst+consensus (+/– reviewer).
- **Label aliasing on/off**.
- **Low-confidence gate** *γ*_*i*_ *< θ* vs. all cells.
- **Two-hop features** on/off; CRF refinement on/off.

## D Datasets and Hyperparameters

STARmap (BZ5, BZ14) use *r* = 700; MERFISH (0.04, 0.09, 0.14, 0.19) use *r* = 600. Default *K* = 3, `smooth_rounds=1–2, num_predict=8` for local models.

## E Results

Results for all six data are shown in Appendix Table 3-8.

**Table 3.** STARmap BZ5 (1500 cells). Metrics: higher is better except CHAOS.

| Method | NMI | HOM | COM | ARI | CHAOS ↓ | PAS | ASW |
| --- | --- | --- | --- | --- | --- | --- | --- |
| baseline_rb | 0.6539 | 0.2488 | 0.4768 | 0.2093 | 0.9604 | 0.1309 | 0.0120 |
| baseline_nearest | 0.6727 | 0.2517 | 0.5067 | 0.2149 | 0.9625 | 0.0281 | 0.0066 |
| baseline_rb_nosmooth | 0.6539 | 0.2488 | 0.4768 | 0.2093 | 0.9548 | 0.1115 | 0.0114 |
| single_llm_allcells | 0.6559 | 0.2505 | 0.4748 | 0.2105 | 0.9604 | 0.1309 | -0.0568 |
| single_llm_lowconf | 0.6539 | 0.2488 | 0.4768 | 0.2093 | 0.9604 | 0.1309 | 0.0120 |
| agentic_2analyst_consensus | 0.6539 | 0.2488 | 0.4768 | 0.2093 | 0.9604 | 0.1309 | 0.0120 |
| agentic_2analyst_consensus_reviewer | 0.6539 | 0.2488 | 0.4768 | 0.2093 | 0.9604 | 0.1309 | 0.0120 |

**Table 4.** STARmap BZ14 (1500 cells). Metrics: higher is better except CHAOS.

| Method | NMI | HOM | COM | ARI | CHAOS ↓ | PAS | ASW |
| --- | --- | --- | --- | --- | --- | --- | --- |
| baseline_rb | 0.4126 | 0.1654 | 0.2742 | 0.2268 | 0.9467 | -0.0207 | -0.0174 |
| baseline_nearest | 0.6153 | 0.2459 | 0.4108 | 0.2900 | 0.9502 | -0.0280 | -0.0179 |
| baseline_rb_nosmooth | 0.4335 | 0.1739 | 0.2876 | 0.2330 | 0.9412 | -0.0159 | -0.0151 |
| single_llm_allcells | 0.4158 | 0.1662 | 0.2777 | 0.2265 | 0.9479 | -0.0291 | -0.0205 |
| single_llm_lowconf | 0.4183 | 0.1672 | 0.2791 | 0.2280 | 0.9474 | -0.0295 | -0.0201 |
| agentic_2analyst_consensus | 0.4521 | 0.1807 | 0.3017 | 0.2444 | 0.9483 | -0.0281 | -0.0198 |
| agentic_2analyst_consensus_reviewer | 0.4521 | 0.1807 | 0.3017 | 0.2444 | 0.9483 | -0.0281 | -0.0198 |

**Table 5.** MERFISH 0.04 (1500 cells). Metrics: higher is better except CHAOS.

| Method | NMI | HOM | COM | ARI | CHAOS ↓ | PAS | ASW |
| --- | --- | --- | --- | --- | --- | --- | --- |
| baseline_rb | 0.4203 | 0.1260 | 0.6320 | 0.0552 | 0.9080 | -0.0247 | 0.0202 |
| baseline_nearest | 0.3980 | 0.1326 | 0.3990 | 0.0463 | 0.7420 | -0.0374 | 0.0112 |
| baseline_rb_nosmooth | 0.3723 | 0.1110 | 0.5771 | 0.0476 | 0.9002 | -0.0383 | 0.0135 |
| single_llm_allcells | 0.2009 | 0.0561 | 0.4808 | 0.0141 | 0.9214 | -0.0681 | -0.0100 |
| single_llm_lowconf | 0.2949 | 0.0853 | 0.5418 | 0.0290 | 0.9061 | -0.0549 | 0.0143 |
| agentic_2analyst_consensus | 0.3263 | 0.0955 | 0.5595 | 0.0349 | 0.9045 | -0.0529 | 0.0113 |
| agentic_2analyst_consensus_reviewer | 0.3263 | 0.0955 | 0.5595 | 0.0349 | 0.9045 | -0.0529 | 0.0113 |

**Table 6.** MERFISH 0.09 (1500 cells). Metrics: higher is better except CHAOS.

| Method | NMI | HOM | COM | ARI | CHAOS ↓ | PAS | ASW |
| --- | --- | --- | --- | --- | --- | --- | --- |
| baseline_rb | 0.3483 | 0.1012 | 0.6256 | 0.0365 | 0.9015 | 0.0197 | -0.0169 |
| baseline_nearest | 0.5964 | 0.2050 | 0.5466 | 0.0969 | 0.7217 | 0.1220 | -0.0123 |
| baseline_rb_nosmooth | 0.6675 | 0.2332 | 0.5870 | 0.1235 | 0.7060 | 0.1692 | -0.0005 |
| single_llm_allcells | 0.0689 | 0.0180 | 0.4253 | 0.0014 | 0.9731 | -0.0801 | -0.0817 |
| single_llm_lowconf | 0.1851 | 0.0504 | 0.5600 | 0.0096 | 0.9390 | -0.0673 | -0.0042 |
| agentic_2analyst_consensus | 0.1895 | 0.0522 | 0.5101 | 0.0113 | 0.9312 | -0.0662 | 0.0005 |
| agentic_2analyst_consensus_reviewer | 0.1895 | 0.0522 | 0.5101 | 0.0113 | 0.9312 | -0.0662 | 0.0005 |

**Table 7.** MERFISH 0.14 (1500 cells). Metrics: higher is better except CHAOS.

| Method | NMI | HOM | COM | ARI | CHAOS ↓ | PAS | ASW |
| --- | --- | --- | --- | --- | --- | --- | --- |
| baseline_rb | 0.9637 | 0.3608 | 0.7251 | 0.2783 | 0.7916 | 0.1679 | 0.0143 |
| baseline_nearest | 0.8052 | 0.3068 | 0.5855 | 0.2336 | 0.6120 | 0.1518 | 0.0081 |
| baseline_rb_nosmooth | 0.7043 | 0.2666 | 0.5185 | 0.1893 | 0.6273 | 0.1019 | 0.0118 |
| single_llm_allcells | 0.8351 | 0.2792 | 0.8275 | 0.2103 | 0.9390 | 0.0188 | 0.0152 |
| single_llm_lowconf | 0.8348 | 0.2778 | 0.8391 | 0.2105 | 0.9418 | 0.0177 | 0.0304 |
| agentic_2analyst_consensus | 0.8502 | 0.2843 | 0.8422 | 0.2126 | 0.9415 | 0.0250 | 0.0153 |
| agentic_2analyst_consensus_reviewer | 0.8502 | 0.2843 | 0.8422 | 0.2126 | 0.9415 | 0.0250 | 0.0153 |

**Table 8.** MERFISH 0.19 (1500 cells). Metrics: higher is better except CHAOS.

| Method | NMI | HOM | COM | ARI | CHAOS ↓ | PAS | ASW |
| --- | --- | --- | --- | --- | --- | --- | --- |
| baseline_rb | 0.6912 | 0.2432 | 0.5967 | 0.1672 | 0.8328 | -0.0854 | -0.0117 |
| baseline_nearest | 0.7371 | 0.2702 | 0.5794 | 0.1766 | 0.6376 | 0.2677 | 0.0093 |
| baseline_rb_nosmooth | 0.7723 | 0.2862 | 0.5936 | 0.2002 | 0.6363 | 0.2423 | 0.0079 |
| single_llm_allcells | 0.5460 | 0.1694 | 0.7027 | 0.0881 | 0.8232 | 0.2694 | 0.0506 |
| single_llm_lowconf | 0.2147 | 0.0598 | 0.5250 | 0.0098 | 0.9732 | -0.0070 | -0.0534 |
| agentic_2analyst_consensus | 0.6680 | 0.2238 | 0.6574 | 0.1499 | 0.8226 | 0.0622 | -0.0638 |
| agentic_2analyst_consensus_reviewer | 0.6680 | 0.2238 | 0.6574 | 0.1499 | 0.8226 | 0.0622 | -0.0638 |

### Failure Modes

Despite improved spatial coherence, lightweight LLM agents occasionally exhibited mode collapse, assigning a dominant label across all regions, particularly in low-signal MER-FISH subsets or when confidence margins were small. We attribute this to limited prompt diversity and weak separation among prototype features, which can cause deterministic bias in consensus voting. In addition, some free-tier models failed to return valid JSON outputs under multi-agent prompting, requiring fallback to rule-based predictions. These issues highlight the sensitivity of agentic reasoning to both prompt structure and contextual grounding, suggesting the need for adaptive temperature control, per-label calibration, and hybrid symbolic–neural integration in future iterations.

### Additional Details

The appendix includes extended implementation notes, ablation results, and dataset statistics for all six spatial transcriptomics datasets (two STARmap and four MERFISH samples). We report parameter settings for neighborhood radius, prototype cardinality, and LLM temperature, along with full per-model metrics for each baseline (rule-based, nearest-prototype, no-smoothing, and agentic variants). Runtime analysis on 500–1500 cells demonstrates that lightweight models (`llama3.2, qwen3`) can perform full annotation within minutes on a single GPU-free node. We also include qualitative visualization of predicted niche boundaries and pairwise label coherence maps, confirming that agentic models improve spatial smoothness without compromising label diversity. All scripts and evaluation logs will be released to support reproducibility.

## References

Quentin Blampey, Kevin Mulder, Margaux Gardet, Stergios Christodoulidis, Charles-Antoine Dutertre, Fabrice André, Florent Ginhoux, and Paul-Henry Cournède. 2024. Sopa: a technology-invariant pipeline for analyses of image-based spatial omics. Nature Communications, 15(1):4981.

Kok Hao Chen, Alistair N Boettiger, Jeffrey R Moffitt, Siyuan Wang, and Xiaowei Zhuang. 2015. Spatially resolved, highly multiplexed rna profiling in single cells. Science, 348(6233):aaa6090.

Yiqun Chen and James Zou. 2024. Genept: a simple but effective foundation model for genes and cells built from chatgpt. bioRxiv, pages 2023–10.

Haotian Cui, Chloe Wang, Hassaan Maan, Kuan Pang, Fengning Luo, Nan Duan, and Bo Wang. 2024. scgpt: toward building a foundation model for single-cell multi-omics using generative ai. Nature methods, 21(8):1470–1480.

Kangning Dong and Shihua Zhang. 2022. Deciphering spatial domains from spatially resolved transcriptomics with an adaptive graph attention auto-encoder. Nature communications, 13(1):1739.

Ruben Dries, Qian Zhu, Rui Dong, Chee-Huat Linus Eng, Huipeng Li, Kan Liu, Yuntian Fu, Tianxiao Zhao, Arpan Sarkar, Feng Bao, and 1 others. 2021. Giotto: a toolbox for integrative analysis and visualization of spatial expression data. Genome biology, 22(1):78.

Minsheng Hao, Jing Gong, Xin Zeng, Chiming Liu, Yucheng Guo, Xingyi Cheng, Taifeng Wang, Jianzhu Ma, Xuegong Zhang, and Le Song. 2024. Largescale foundation model on single-cell transcriptomics. Nature methods, 21(8):1481–1491.

Jian Hu, Xiangjie Li, Kyle Coleman, Amelia Schroeder, Nan Ma, David J Irwin, Edward B Lee, Russell T Shinohara, and Mingyao Li. 2021. Spagcn: Integrating gene expression, spatial location and histology to identify spatial domains and spatially variable genes by graph convolutional network. Nature methods, 18(11):1342–1351.

Yahui Long, Kok Siong Ang, Mengwei Li, Kian Long Kelvin Chong, Raman Sethi, Chengwei Zhong, Hang Xu, Zhiwei Ong, Karishma Sachaphibulkij, Ao Chen, and 1 others. 2023. Spatially informed clustering, integration, and deconvolution of spatial transcriptomics with graphst. Nature Communications, 14(1):1155.

Valentine Svensson, Sarah A Teichmann, and Oliver Stegle. 2018. Spatialde: identification of spatially variable genes. Nature methods, 15(5):343–346.

Xiao Wang, William E Allen, Matthew A Wright, Emily L Sylwestrak, Nikolay Samusik, Sam Vesuna, Kathryn Evans, Cindy Liu, Charu Ramakrishnan, Jia Liu, and 1 others. 2018a. Three-dimensional intacttissue sequencing of single-cell transcriptional states. Science, 361(6400):eaat5691.

Xiao Wang, William E Allen, Matthew A Wright, Emily L Sylwestrak, Nikolay Samusik, Sam Vesuna, Kathryn Evans, Cindy Liu, Charu Ramakrishnan, Jia Liu, and 1 others. 2018b. Three-dimensional intacttissue sequencing of single-cell transcriptional states. Science, 361(6400):eaat5691.

Huanhuan Wei, Xiao Luo, Hongyi Yu, Jinping Liang, Luning Yang, Lixing Lin, Alexandra Popa, and Xiting Yan. 2025. Identifying cellular niches in spatial transcriptomics: An investigation into the capabilities of large language models. In Proceedings of the 63rd Annual Meeting of the Association for Computational Linguistics (Volume 1: Long Papers), pages 9275–9289.

Hongzhi Wen, Wenzhuo Tang, Wei Jin, Jiayuan Ding, Renming Liu, Xinnan Dai, Feng Shi, Lulu Shang, Hui Liu, and Yuying Xie. 2023. Single cells are spatial tokens: Transformers for spatial transcriptomic data imputation. arXiv preprint arXiv:2302.03038.

Chang Xu, Xiyun Jin, Songren Wei, Pingping Wang, Meng Luo, Zhaochun Xu, Wenyi Yang, Yideng Cai, Lixing Xiao, Xiaoyu Lin, and 1 others. 2022. Deepst: identifying spatial domains in spatial transcriptomics by deep learning. Nucleic Acids Research, 50(22):e131–e131.

Zhiyuan Yuan, Fangyuan Zhao, Senlin Lin, Yu Zhao, Jianhua Yao, Yan Cui, Xiao-Yong Zhang, and Yi Zhao. 2024. Benchmarking spatial clustering methods with spatially resolved transcriptomics data. Nature Methods, 21(4):712–722.

Meng Zhang, Stephen W Eichhorn, Brian Zingg, Zizhen Yao, Kaelan Cotter, Hongkui Zeng, Hongwei Dong, and Xiaowei Zhuang. 2021. Spatially resolved cell atlas of the mouse primary motor cortex by merfish. Nature, 598(7879):137–143.

Chongyue Zhao, Zhongli Xu, Xinjun Wang, Shiyue Tao, William A MacDonald, Kun He, Amanda C Poholek, Kong Chen, Heng Huang, and Wei Chen. 2024. Innovative super-resolution in spatial transcriptomics: a transformer model exploiting histology images and spatial gene expression. Briefings in Bioinformatics, 25(2):bbae052.

Edward Zhao, Matthew R Stone, Xing Ren, Thomas Pulliam, Paul Nghiem, Jason H Bielas, and Raphael Gottardo. 2020. Bayesspace enables the robust characterization of spatial gene expression architecture in tissue sections at increased resolution. bioRxiv, pages 2020–09.

